# Inhibition of cell membrane ingression at the division site by cell wall in fission yeast

**DOI:** 10.1101/2020.04.06.027193

**Authors:** Ting Gang Chew, Tzer Chyn Lim, Yumi Osaki, Junqi Huang, Anton Kamnev, Masako Osumi, Mohan K. Balasubramanian

**Author notes:** Co-first authors (Ting Gang Chew, Tzer Chyn Lim). Correspondence to: Mohan K. Balasubramanian.

## Abstract

Eukaryotic cells assemble an actomyosin ring during cytokinesis to function as a force-generating machine to drive membrane invagination, and to counteract the intracellular pressure and the cell surface tension. It is unclear whether additional factors such as the extracellular matrix (cell wall in yeasts and fungi) affect the actomyosin ring contraction. While studying the fission yeast β-glucan synthase mutant *cps1*-191, which is defective in division septum synthesis and actomyosin ring contraction, we found that significantly weakening of the extracellular glycan matrix caused the spheroplasts to divide at the non-permissive condition. This division was dependent on a functional actomyosin ring and vesicular trafficking, but independent of normal septum synthesis. *cps1*-191 cells with weakened extracellular glycan matrix divide non-medially with a much slower ring contraction rate compared to wild type cells under similar conditions, which we term as cytofission. Interestingly, the high turgor pressure appears to play minimal roles in inhibiting ring contraction in *cps1*-191 mutants as decreasing the turgor pressure alone does not enable cytofission. We propose that during cytokinesis, the extracellular glycan matrix restricts actomyosin ring contraction and membrane ingression, and remodeling of the extracellular components through division septum synthesis relieves the inhibition and facilitates actomyosin ring contraction.

## Introduction

Animal cells and yeast require assembly and contraction of an actomyosin ring during cytokinesis. In fission yeast, the actomyosin ring contracts to drive membrane ingression, and coordinates with the septum assembly machinery to deposit cell wall materials at the division site (Ramos et al., 2019). The septum assembly machinery consists of glucan synthases such as β-glucan synthase Bgs1/Cps1 and α-glucan synthase Ags1/Mok1. Cps1 synthesizes the β-glucan matrix at the division site and couples the extracellular glycan matrix to the actomyosin ring via intermediate protein complexes (Sethi et al., 2016; Davidson et al., 2016). The deposition of extracellular glycan matrix coordinates with actomyosin ring contraction and stabilizes the contracting actomyosin ring at the division site (Muñoz et al., 2013).

Whether the extracellular glycan matrix has any influence on the actomyosin ring contraction apart from its roles in ring stability during cytokinesis has not been examined closely. In this study, we used the thermosensitive allele of β-1,3-glucan synthase, *cps1*-191 to address this question. The *cps1*-191 mutant is defective in β-glucan synthesis and arrests with a non-contracting actomyosin ring at the non-permissive temperature. Interestingly, we found that weakening of the extracellular glycan matrix in *cps1*-191 mutant at the non-permissive temperature has enabled actomyosin ring contraction and membrane ingression.

## Results and discussion

Under non-permissive temperature, the β-glucan synthase mutant *cps1*-191 assembles actomyosin rings that do not contract (Liu et al., 2000). It has been suggested that β-glucan synthesis at the division site is required to overcome the high intracellular turgor pressure during cytokinesis, and that the actomyosin ring may not be able to overcome the high turgor (Proctor et al., 2012). To test if the turgor pressure inhibited ring contraction in *cps1*-191 mutants, we cultured *cps1*-191 cells in medium containing 0.8 M sorbitol to decrease the turgor pressure to that inside the cells, and we added 2-deoxyglucose (2-DG) to this culture to prevent further glucan synthesis at the division site and elsewhere in the cell (Megnet, 1965; Svoboda and Smith, 1972; Osumi et al., 1998). As previously reported, actomyosin rings of *cps1*-191 cells maintained in normal turgor pressure did not undergo contraction (Figure 1A; movie 1). We occasionally observed that parts of the *cps1*-191 cells swelled into a bump and the cells lysed eventually with a collapsing ring (Figure 1B; movie 2). Culturing *cps1*-191 cells in medium containing 0.8 M sorbitol, did not increase actomyosin ring contraction events and phenotypically these cells resembled *cps1*-191 grown under normal growth conditions, in which a high intracellular turgor pressure is maintained (Figure 1C - movie 3 and 1D - movie 4). Thus, our results showed that a decreased turgor pressure does not allow ring contraction in *cps1*-191 mutants.

**Figure 1.**
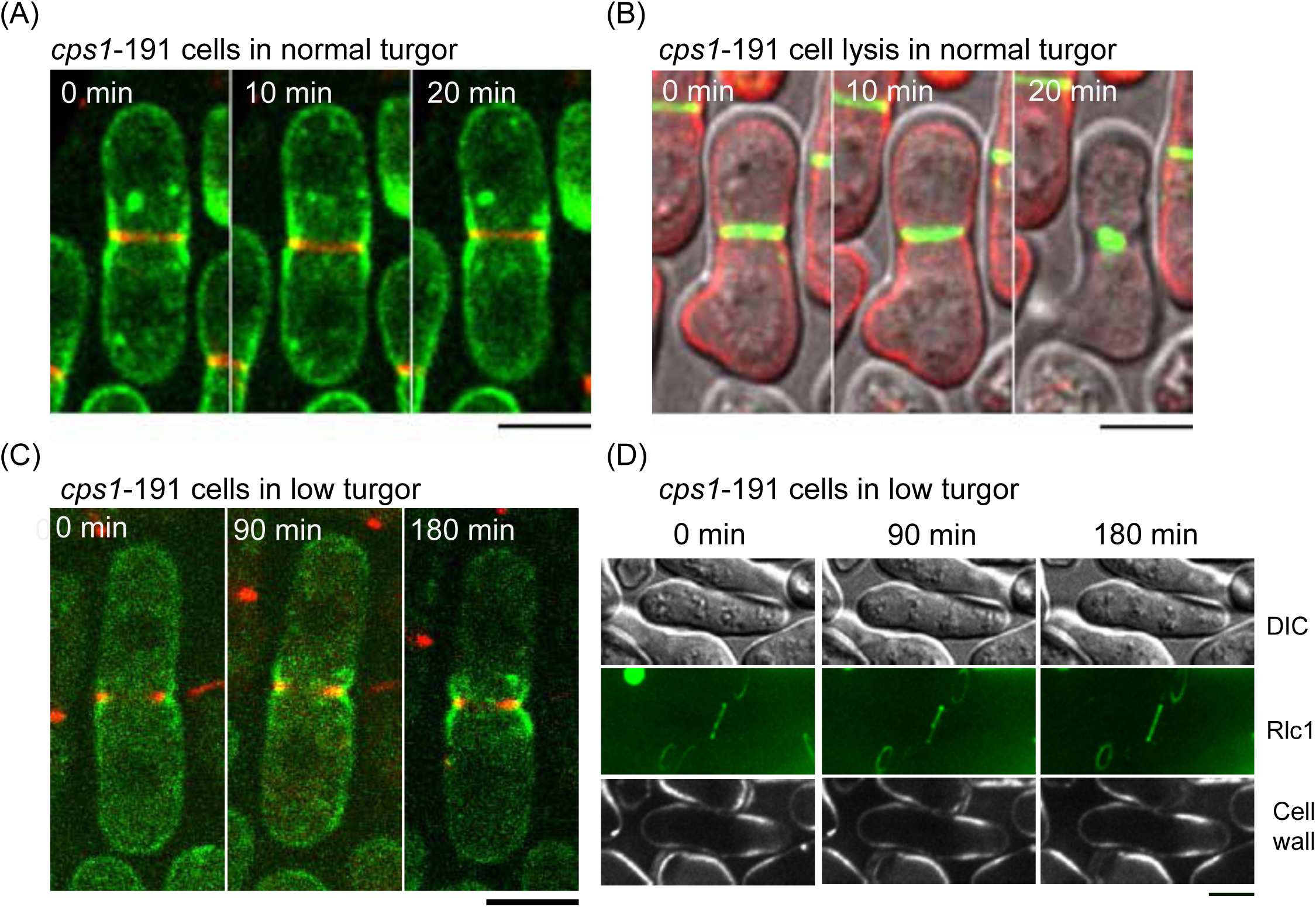
Lowering down turgor pressure does not allow cell membrane ingression in *cps1* mutant cells. (A) The *cps1*-191 GFP-*psy1 rlc1*-tdTomato cells were cultured at 36°C for 6.5 hours and continued to image at 36°C. Green: GFP-*psy1*. Red: *rlc1*-tdTomato. (B) Some *cps1*-191 GFP-*psy1 rlc1*-tdTomato cells lysed after more than 6.5 hours of incubation at the restrictive temperature. Green: *rlc1*-tdTomato. Red: GFP-*psy1*. (C) The *cps1*-191 GFP-*psy1 rlc1*-tdTomato cells were cultured at the restrictive temperature for 6.5 hours and continued to image at 36°C in medium containing 0.8 M sorbitol to lower down the turgor pressure. Green: GFP-*psy1*. Red: *rlc1*-tdTomato. (D) The *cps1*-191 GFP-*psy1 rlc1*-tdTomato cells were cultured at the restrictive temperature for 6.5 hours and continued to image at 36°C in medium containing 0.8 M sorbitol. Calcofluor was used to stain the cell wall. Scale bars: 5 μm

Next, we considered the possibility that the extracellular glycan matrix inhibited ring contraction and membrane ingression in *cps1*-191 mutants. The Cps1 is a transmembrane protein that (along with other integral membrane proteins, such as Ags1, Bgs3, and Bgs4) links actomyosin rings underneath the cell membrane to the extracellular glycan matrix (Muñoz et al., 2013; Cortés et al., 2012; Davidson et al., 2016; Sethi et al., 2016; Arasada and Pollard, 2015; Cortés et al., 2005). It was possible that in the absence of division septum synthesis (and thereby cell wall remodeling), the actomyosin rings are permanently fixed to the inactive *cps1*-191 gene-product anchored at the division site. To test if this was the case, we weakened the extracellular glycan matrix by treating the *cps1*-191 cells with cell wall lytic enzymes and further blocking new cell wall and septum synthesis by supplementing the culture with 2-DG. Interestingly, upon weakening of the cell wall, myosin rings in *cps1*-191 mutant expressing the regulatory light chain of myosin tagged with the fluorescent protein tdTomato (Rlc1-tdTomato) underwent contraction coupled with membrane ingression (Figure 2A; GFP-tagged Syntaxin-like protein Psy1 was used as a cell membrane marker; n = 19 spheroplasts; movies 5 and 6). Consistently, contracting actin rings labeled with the Lifeact-mCherry were also detected in the *cps1*-191 mutant upon weakening of cell wall, suggesting that the actomyosin rings were driving the contraction and membrane ingression (Figure 2B; n = 5 spheroplasts; movies 7 and 8). These mutant spheroplasts with weakened cell wall often divided non-medially into two, and the rings contracted at much reduced rate (0.061 ± 0.212 μm/min, n_spheroplast_ = 8) compared to wild-type cells (Figure 2B; 0.299 ± 0.059 μm/min, n_cell_ = 14,) (Figure 2C). We frequently observed that the rings contracted till mid-phase of division and disassembled before completion of cytokinesis. The spheroplasts however went on to divide into two entities (Figure 2A and 2B). The segregation of daughter nuclei in the *cps1*-191 spheroplasts was often not coordinated with the cytokinesis, with some spheroplasts have multiple nuclei in one of the daughter entities or have cleaved nuclei, presumably due to the non-medial division (Supplementary figure 1; movies 9-11). Since the *cps1*-191 mutant spheroplast division was morphologically different from normal fission yeast cell division and was mimicking the morphological changes of some animal cells during division, we have called this type of division as cytofission.

**Figure 2.**
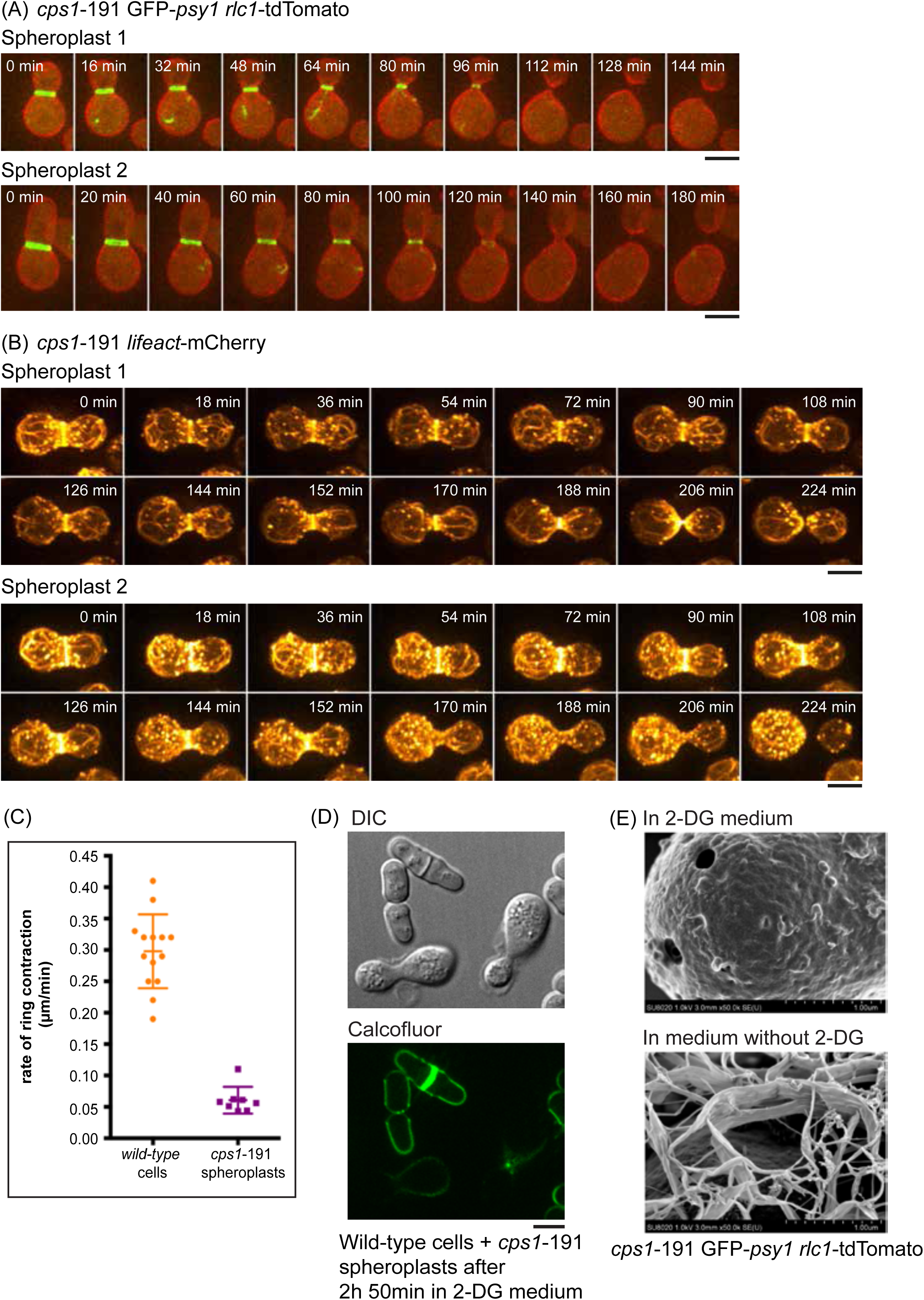
Weakening of cell wall allows ring contraction and cell membrane ingression. (A) Two examples of *cps1*-191 spheroplasts underwent cytofission at 36°C. The *cps1*-191 GFP-*psy1 rlc1*-tdTomato cells were cultured at 36°C for 6.5 hours, processed into spheroplasts, and recovered for 1 hour at 36°C prior to imaging. (B) Two examples of *cps1*-191 spheroplasts expressing Lifeact-mCherry underwent cytofission at 36°C. The *cps1*-191 GFP-*psy1 lifeact*-mCherry cells were cultured at 36°C for 6.5 hours, processed into spheroplasts, and recovered for 1 hour at 36°C prior to imaging. (C) Quantification of the rate of ring contraction in wild type cells and *cps1*-191 spheroplasts undergoing cytofission. (D) The wild type cells and *cps1*-191 GFP-*psy1 rlc1*-tdTomato spheroplasts were stained with the calcofluor dye. (E) Electron micrographs of *cps1*-191 GFP-*psy1 rlc1*-tdTomato spheroplasts regenerated in medium with or without 2-DG. Scale bars: 5 μm Error bars: standard deviation

Analysis of the extracellular glycan matrix using calcofluor staining (a division septum-specific fluorochrome) (G. Cortés et al., 2018) in cells undergoing cytofission revealed that the division site of *cps1*-191 spheroplasts undergoing cytofission contained significantly reduced β-glucan materials (Figure 2D). Further study with the high-resolution scanning electron microscopy showed that the glucan fibrils regenerated in *cps1*-191 spheroplasts without 2-DG (Figure 2E; bottom panel) while the fibrils were not noticeable in *cps1*-191 spheroplasts with 2-DG (Figure 2E; top panel). The glucan fibrils commonly present at the division site of fission yeast was largely absent in *cps1*-191 spheroplasts undergoing cytofission (Supplementary figure 2). Taken together, we showed that weakening of cell wall in *cps1*-191 cells at non-permissive temperature facilitates a novel cytofission event that leads to division of one spheroplast into two in the absence of detectable division-septum growth. Our results also suggested that the extracellular glycan matrix anchored to the actomyosin rings negatively regulates the ring contraction and membrane ingression.

A reduction of β-glucan may result in an increased amount of α-glucan in the cell wall of fission yeast. To test if the cytofission of *cps1*-191 spheroplasts were due to the synthesis of α-glucan at the division site, we prepared the *cps1*-191 *mok1*-664 double mutant spheroplasts containing the thermosensitive alleles of both α- and β-glucan synthases, and imaged the myosin rings and cell membrane in this double mutant spheroplasts at the non-permissive temperature. Similar to the *cps1*-191 spheroplast, the *cps1*-191 *mok1*-664 double mutant spheroplasts underwent cytofission (Figure 3A, n = 8; movie 12), suggesting that α-glucan and β-glucan synthesis did not contribute significantly to the cytofission events.

**Figure 3.**
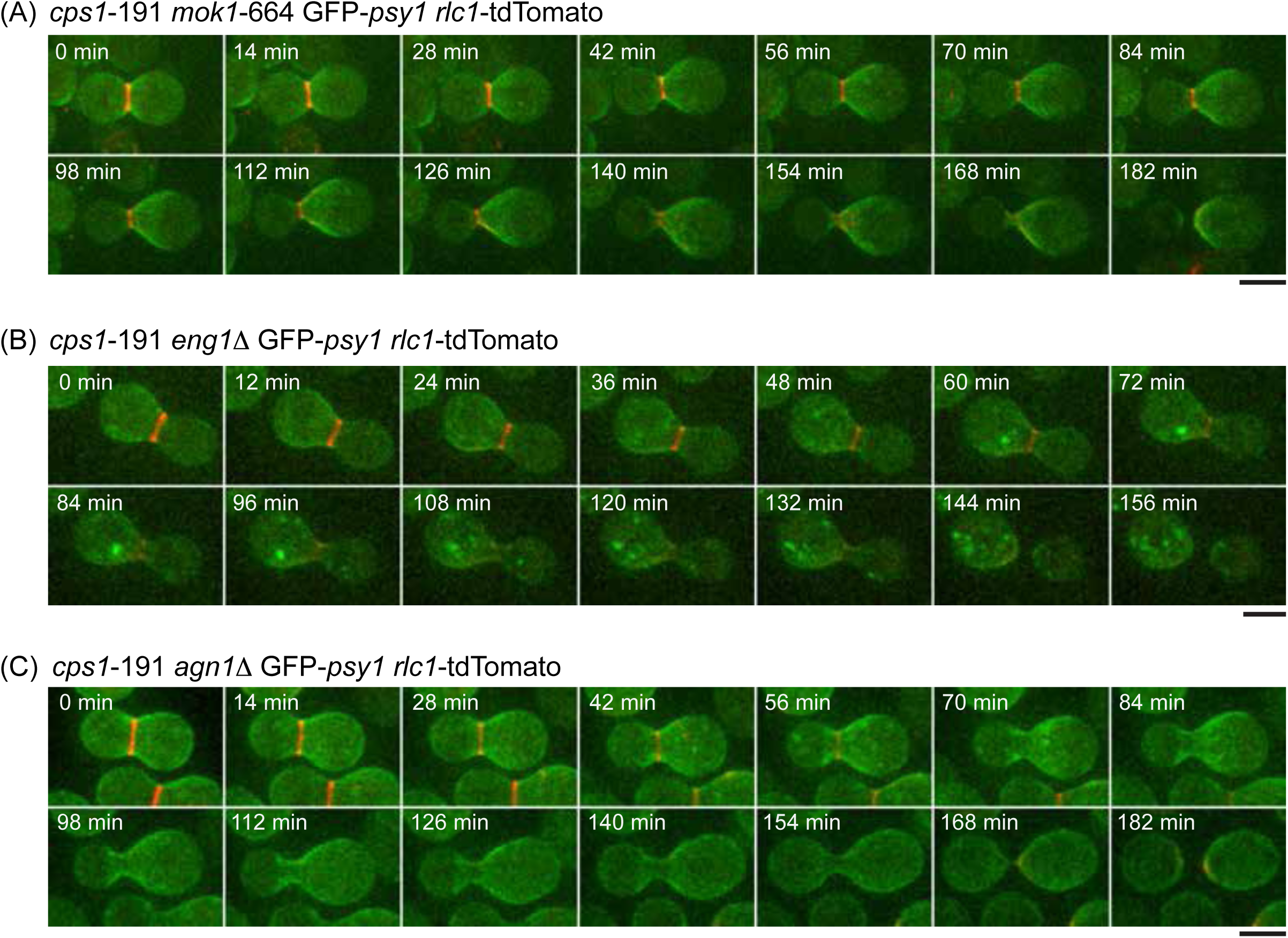
Cps1 mutant spheroplasts undergo cytofission independent of the α-glucan synthase and exoglucanases. (A) An example of cps1-191 *mok1*-664 GFP-*psy1 rlc1*-tdTomato spheroplasts underwent cytofission. (B) An example of *cps1*-191 *agn1*Δ GFP-*psy1 rlc1*-tdTomato spheroplasts underwent cytofission. (C) An example of *cps1*-191 *eng1*Δ GFP-*psy1 rlc1*-tdTomato spheroplasts underwent cytofission. Scale bars: 5 μm

Normal fission yeast cells that just complete ring contraction and membrane ingression are not entirely separated until the digestion of division septum connecting the two newly-divided cells (Sipiczki, 2007). This process is achieved in fission yeast through the action of exoglucanases (Martín-Cuadrado et al., 2003; Dekker et al., 2004; García et al., 2005). We tested if proteins involved in the separation of fission yeast cells were also involved in the cytofission, which would be expected if trace amounts of division septum had been deposited during ring contraction. To this end, we constructed double mutant spheroplasts of *cps1*-191 lacking the exoglucanases *eng1* (β-glucanase) and *agn1* (α-glucanase), respectively. Similar to the single mutant *cps1*-191, the double mutants lacking either of the two exoglucanases underwent cytofission upon weakening of the cell wall (Figure 3B and 3C; movies 13 and 14). The results indicated that the cytofission events of *cps1*-191 mutants does not require the break-down of cell wall materials by exoglucanases, even though cytofission leads to the complete separation of spheroplasts.

In ∼70% of the *cps1*-191 spheroplasts (37 out of 53 spheroplasts) that underwent cytofission, the rings contracted till mid-phase of division and disassembled before division into two entities. We tested if the ESCRT abscission complex was involved in the cytofission by removing two of the ESCRT proteins Vps4 and Vps20 in the *cps1*-191 spheroplasts. The *cps1*-191 *vps4*Δ and *cps1*-191 *vps20*Δ double mutant spheroplasts underwent cytofission like in the single *cps1*-191 mutant spheroplast (Supplementary figure 3; movies 15 and 16). It is possible that the completion of cytofission without the actomyosin rings was achieved via an unknown cell abscission mechanism.

Previous studies suggested under certain circumstances, some eukaryotic cells are able to divide without an actomyosin ring (Flor-Parra et al., 2014; Proctor et al., 2012; Dix et al., 2018; Choudhary et al., 2013). To see if the cytofission was driven by contraction of the actomyosin ring, we first perturbed the functions of rings using Latrunculin-A (Lat-A) to inhibit actin polymerization (Fujiwara et al., 2018; Morton et al., 2000). *cps1*-191 cells treated with DMSO underwent ring contraction, membrane ingression, and completed cytofission (Figure 4A; movie 17; n = 7 spheroplasts). By contrast, *cps1*-191 cells treated with Lat-A underwent ring disassembly and failed to divide into two entities. Interestingly, the smaller entity retracted into the bigger entity, probably due to the imbalance of intracellular pressures (Figure 4B; movie 18; n = 8 spheroplasts).

**Figure 4.**
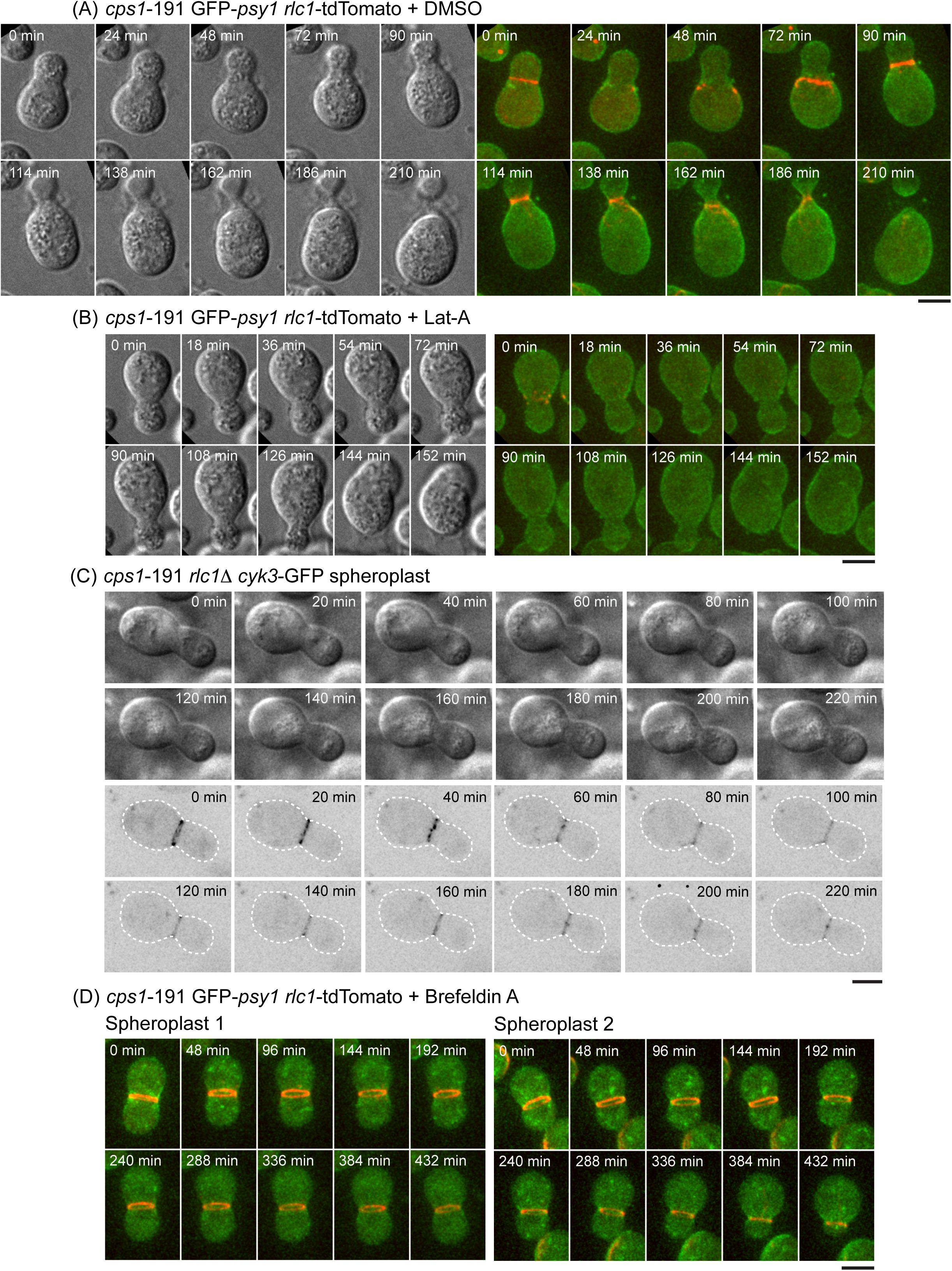
The function of actomyosin rings is required in the Cps1 mutant spheroplasts to undergo cytofission. (A) The *cps1*-191 GFP-*psy1 rlc1*-tdTomato spheroplasts underwent cytofission in the presence of DMSO. Left panel shows the DIC images; right panel shows the fluorescence micrographs. (B) The *cps1*-191 GFP-*psy1 rlc1*-tdTomato spheroplasts were incubated with 150 μm Lat-A. Left panel shows the DIC images; right panel shows the fluorescence micrographs. (C) The *rlc1*Δ *cyk3*-GFP spheroplasts failed to undergo cytofission at 36°C. The *rlc1*Δ *cyk3*-GFP cells were cultured at 36°C for 6.5 hours, processed into spheroplasts, and then recovered in minimal medium containing sorbitol prior to imaging at 36°C. Top panel shows the DIC images; bottom panel shows the fluorescence micrographs. (D) The *cps1*-191 GFP-*psy1 rlc1*-tdTomato spheroplasts failed to undergo cytofission in the presence of 75 μM brefeldin A. Scale bars: 5 μm

Next, we perturbed the myosin component of actomyosin rings by deleting *rlc1*, the regulatory light chain of myosin II (Naqvi et al., 2000; Le Goff et al., 2000; Pollard et al., 2017), in *cps1*-191 mutants. It has been shown that the cells lacking *rlc1* (*rlc1*Δ) fail to undergo cytokinesis at low temperature, but at high temperatures, the *rlc1*Δ cells assemble an intact actomyosin ring that contracts normally (Naqvi et al., 2000) (Supplementary figure 4; movies 19 and 20; n = 23 cells). We used this differential temperature requirement to test the essentiality of actomyosin ring functions in cytofission. If the actomyosin ring was essential in driving cytofission, the absence of *rlc1* might render the cells with weakened cell wall unable to undergo cytofission at the high temperature, which is normally permissive for *rlc1*Δ cells alone (Naqvi et al., 2000). Consistent with the LatA experimental findings, the double mutant *cps1*-191 *rlc1*Δ with weakened cell wall did not undergo ring contraction at the restrictive temperature (Figure 4C, movies 21 and 22; n = 28 spheroplasts), whereas the single mutant of *cps1*-191 could undergo cytofission.

Next, we tested if targeted membrane deposition at the division site facilitates actomyosin ring contraction in cytofission. When the vesicular trafficking across the endomembrane system was inhibited using brefeldin A in the *cps1*-191 spheroplasts, the myosin rings were not able to contract to drive cytofission events (Figure 4D; Movies 23 and 24). This result suggested that addition of cell membrane via targeted membrane trafficking at the division site is required to enable cytofission.

Our study reveals that the extracellular glycan matrix inhibits actomyosin ring contraction in the absence of cell wall remodeling and division septum synthesis. When the inhibition is relieved by experimental treatments like ones reported in this study, or by septum synthesis, the actomyosin ring contracts to drive the membrane ingression. A previous study by Proctor *et al*. analyzed *cps1*-191 mutants and explained that the failure of membrane ingression in the mutant was due to a defect in division-septum assembly (Proctor et al., 2012). The authors also proposed that the high intracellular turgor pressure prevents actomyosin ring contraction in fission yeast (Proctor et al., 2012). We tested this model by lowering the turgor pressure in *cps1*-191 mutant cells and found that it was not sufficient to enable membrane ingression in *cps1*-191 cells. However, the ability of *cps1*-191 mutant cells to divide upon weakening of cell wall indicates that the actomyosin ring in *cps1*-191 mutant cells is capable of driving membrane ingression even when the division septum assembly is defective. The fact that ring contraction is slower during cytofission however, agrees better with the work of O’ Shaughnessy and colleagues, who have proposed that the rate of septum synthesis sets the rate of cytokinesis (Stachowiak et al., 2014). It is possible that our work reveals the highest rate of actomyosin ring contraction when confronted with membrane drag and viscous drag of the cytosol. In fission yeast spheroplasts, the actomyosin ring is probably required at the early phase of cytofission to drive spheroplasts into a dumbbell shape with high curvature. The actomyosin ring and other protein complexes like ESCRT are not necessary at the later phase of cytofission when the curvature is sufficiently high to induce spontaneous division. This finding is consistent with a recent study in which the spontaneous curvature in dumbbell-shape lipid vesicles generates constriction forces to induce membrane fission. This leads to the division of a dumbbell-shaped lipid vesicle into two with an increased curvature (Steinkühler et al., 2020).

The yeast cell wall consists of mainly glycan matrix and glycosylated proteins and has been suggested as a functional equivalent of the extracellular matrix (ECM) in animal cells (Muñoz et al., 2013). The mechanical interaction between the cytokinetic actomyosin ring and the ECM is not well understood. A recent study of zebrafish epicardial cells in the heart explants shows the cell-ECM adhesions at the division site. The cell-ECM adhesions lead to the traction forces at the cytokinetic ring that inhibit cytokinesis (Uroz et al., 2019). An early biophysical study also detected a large traction force at the cleavage furrow of the fibroblast cells cultured on an elastic substrate, suggesting an interaction of cytokinetic machinery and ECM (Taylor, 1997). When the cell-ECM adhesion is enhanced during mitosis, the cleavage furrow ingression is inhibited in the epithelial cells (Taneja et al., 2016). Consistently, our study shows that the anchoring of actomyosin rings to the extracellular glycan matrix that do not undergo remodeling (due to a defective Bgs1) prevents the actomyosin ring contraction and cell membrane ingression. Weakening of the extracellular glycan matrix, presumably mimicking a decreased cell-ECM adhesion, has enabled cytofission events.

## Materials and methods

### Yeast strains, medium, and culture conditions

Table S1 lists the *S. pombe* strains used in our study. Standard fission yeast genetic techniques were used to prepare the strains. The rich medium YEA (5 g/l yeast extract, 30 g/l glucose, 225 mg/l adenine) was used to culture cells until mid-log phase at 24°C before the temperature shift. Latrunculin-A (latA) (Enzo Life Sciences; BML-T119) was used at the final concentration of 150 *µ*M to perturb the actin dynamics in spheroplasts. Brefeldin A (Fisher Scientific; 15526276) was used at the final concentration of 75μM to slow down plasma membrane invagination.

### Preparation of *S. pombe* spheroplasts for live-cell imaging

The *cps1*-191 cells used in this study were first cultured in YEA medium at 24°C to mid-log phase (OD_595_ = 0.2-0.5), and then were shifted to 36°C for 6 hours 15 minutes (non-permissive conditions). Twenty milliliters of culture were spun down at 3,000 r.p.m. for 1 minute, and washed once with equal volume of E-buffer (50 mM sodium citrate, 100 mM sodium phosphate, [pH 6.0]). After spinning down the cells and resuspending cells in 5 ml of E-buffer containing 1.2 M sorbitol, the cell suspension was incubated with 30 mg of lysing enzyme (Sigma, L1412) at 36°C with shaking at 80 r.p.m. for 90 minutes. This was followed by continuous incubation with 40 *µ*l of LongLife Zymolyase (G-Biosciences, 1.5 U/*µ*l) at 36°C with shaking at 80 r.p.m. for 60 minutes. After enzymatic digestion, the cell suspensions were spun down at 450 xg for 2 minutes and washed once with 5 ml of E-buffer containing 0.6 M sorbitol. After spinning at 450 xg for 2 minutes, the spheroplasts were recovered in 10 ml EMMA medium (Edinburgh minimal medium with all amino acids and nucleotides supplements) containing 0.8 M sorbitol and 0.5% (v/v) of 1 M 2-deoxyglucose (Sigma, D6134) for 30 minutes at 36°C prior to microscopy imaging.

### Sample preparation for light microscopy

One to two milliliters of spheroplast suspensions were concentrated to 20-100 *µ*l by centrifugation at 450 xg for 2 minutes. About 10 *µ*l of concentrated spheroplasts were loaded onto an Ibidi *µ*-Slide 8-Well glass bottom dish (Cat. No. 80827), and covered with mineral oil (Sigma, M5310) to prevent evaporation during imaging process.

To image cells in Figure 1 and supplementary Figure 4, the *cps1*-191 cells and *rlc1*Δ cells after shifting to non-permissive conditions were treated with buffers used to prepare spheroplasts but with the lysing and lytic enzymes omitted to preserve the cell wall integrity. After the buffer washing, the *cps1*-191 cells in the Figure1 were recovered in EMMA medium with full supplements containing 0.8 M sorbitol and 0.5% 2-DG. For the *rlc1*Δ cells in the supplementary Figure 4, after the buffer washing, the cells were recovered in EMMA medium with full supplements containing 0.8 M sorbitol but not 2-DG to allow septation.

### Sample preparation for electron microscopy

Two hundred and fifty milliliters of cells with OD_595_ 0.2 were collected for spheroplasting. Spheroplasts were prepared with the protoplasting method described above. Spheroplasts were spun down from EMMA with 0.8 M sorbitol and resuspended in phosphate-buffered saline (PBS) with 2.5% glutaraldehyde and 1.2 M sorbitol. Fixation solution was prepared by adding 2% glutaraldehyde and dissolving 1.2 M sorbitol in PBS. The following procedure were done at 4°C and gently (vortex mixer is avoided). After 2 hours incubation at room temperature, spheroplasts were spun down in round bottom tubes. Spheroplasts were resuspended in fixation solution and stood on ice for 2 hours. The spheroplasts were separated into 2 tubes: washed and unwashed samples. Unwashed samples were stored at 4°C. The washed samples were washed with 1 mL PBS containing 1.2 M sorbitol for three times. Lastly the spheroplasts were resuspended in 1 mL PBS containing 1.2 M sorbitol and stored at 4°C before electron microscopy.

For electron microscopy, glutaraldehyde-fixed cells were placed on a slide glass whose surface was pre-treated with 0.1% poly-L-lysine. They were washed with 0.1 M phosphate buffer (pH 7.2), post-fixed with 1% osmium tetroxide at 4°C for 1 hour, dehydrated with graded series of ethanol, and critical point dried with a Leica EM CPD030 apparatus (Leica Microsystems, Vienna). The specimens were coated with osmium tetroxide by osmium coater (Vacuum Device.inc, Japan) and observed with S-3400N and SU8020 scanning electron microscope (Hitachi High Technologies, Tokyo) at 10.0 kV and 1.0 kV respectively (Namiki et al., 2011).

### Light microscopy

The Andor Revolution XD spinning disk confocal microscope was used to image the spheroplasts and cells at 36°C. The microscope was equipped with a Nikon ECLIPSE Ti inverted microscope, Nikon Plan Apo Lambda 100×/1.45N.A. oil immersion objective lens, a spinning-disk system (CSU-X1; Yokogawa), and an Andor iXon Ultra EMCCD camera. The Andor IQ3 software was used to acquire images at the pixel size of 80 nm/pixel or 69 nm/pixel, depending on the camera models. Laser lines at wavelengths of 488 nm or 561 nm were used for the excitation of fluorophores. Most images were acquired with Z-step sizes of 0.2 *µ*m.

### Image analysis

Images were processed using Fiji. The time-lapse montages are maximum intensity projections of Z-stack of specified time points.

## Acknowledgements

We acknowledge members of the M.K. Balasubramanian laboratory, especially Dr. Tomoyuki Hatano, for discussion and excellent suggestions. This work was supported by research grants from Wellcome Trust (WT101885MA) and ERC (GA 671083 - ACTOMYOSIN RING) to MKB.

