## Supplemental Legends for "Inhibition of cell membrane ingression at the division site by cell wall in fission yeast"

**Supplementary Figure Legends**

**Supplementary figure 1.** Nuclear segregation in Cps1 mutant spheroplasts.

(A) Top panel: dividing c*ps1*-191 spheroplasts exhibited normal nucleus distribution to daughter spheroplasts (n = 17). Middle panel: *cps1*-191 spheroplasts showed uneven nucleus segregation in which a daughter nucleus was cleaved (n = 2). Bottom panel: dividing c*ps1*-191 spheroplasts showed uneven nucleus segregation in which both daughter nuclei were in one of the divided spheroplasts (n = 10).

(B) The number of events of nuclear segregation is quantitated and plotted.

Scale bars: 5 μm

**Supplementary figure 2.** Electron micrographs of *cps1*-191 spheroplasts

Electron micrographs of *cps1*-191 GFP-*psy1* *rlc1*-tdTomato spheroplasts regenerated in medium with or without 2-DG.

**Supplementary figure 3.** Separation of *cps1* mutant protoplasts does not require ESCRT

proteins like Vps20 and Vps4.

(A) An example of *cps1*-191 *vps20Δ* GFP-*psy1* *rlc1*-tdTomato protoplasts underwent cytofission.
(B) An example of *cps1*-191 *vps4Δ* GFP-*psy1* *rlc1*-tdTomato protoplasts underwent cytofission.

**Supplementary figure 4.** The *rlc1*Δ *cyk3*-GFP cells underwent cytokinesis at 36°C.

The *rlc1*Δ *cyk3*-GFP cells were cultured at 36°C for 6.5 hours, washed with spheroplasting buffers without lysing and lytic enzymes, and then recovered in minimal medium containing sorbitol prior to imaging at 36°C. Top panel shows the DIC images; bottom panel shows the fluorescence micrographs.

**Supplementary Movies**

Movies 1-24 are linked to the figures in the manuscript and the supplemental material. Please see text and figure legends for more details.
