## SUPPLEMENTAL FIGURES for "Inhibition of cell membrane ingression at the division site by cell wall in fission yeast"

**Supplementary figure 1**

(A) *cps1*-191 *hht*-GFP GFP-*psy1 rlc1*-tdTomato

Normal nucleus distribution

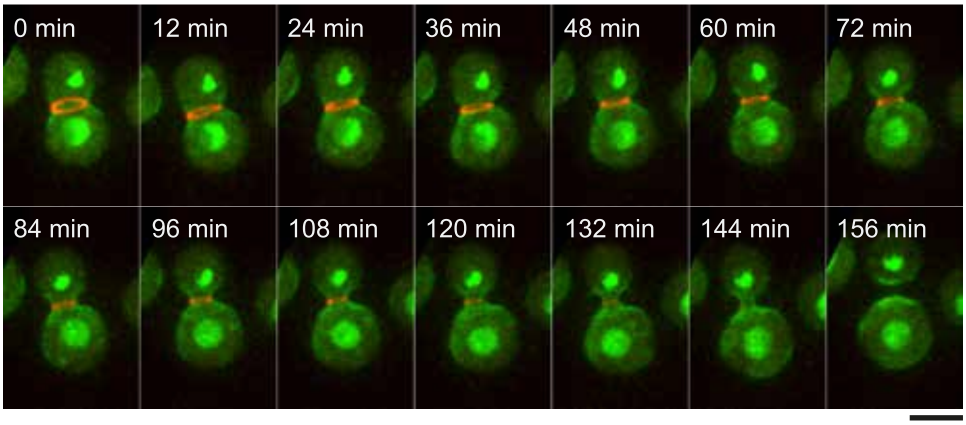


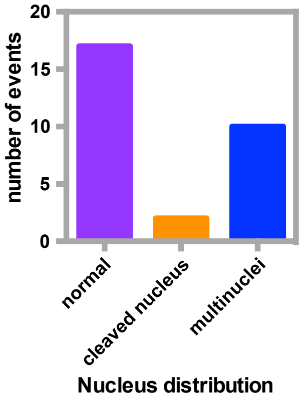


Cleaved nucleus

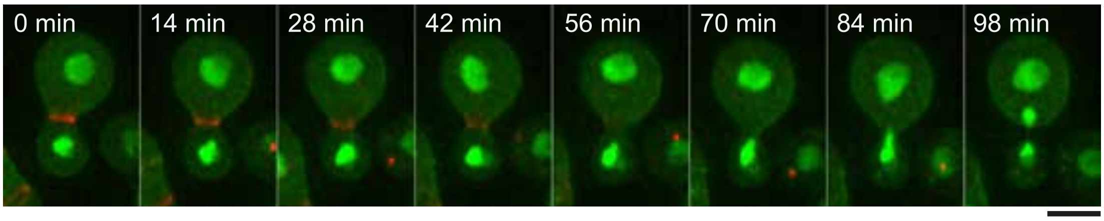


Multinuclei

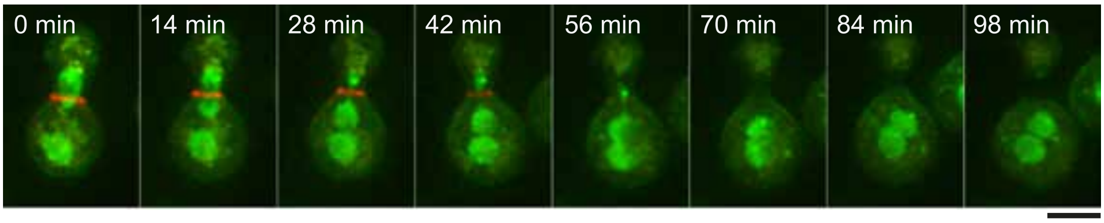

**Supplementary figure 2**

*cps1*-191 GFP-*psy1* *rlc1*-tdTomato

In 2-DG medium    In medium without 2-DG

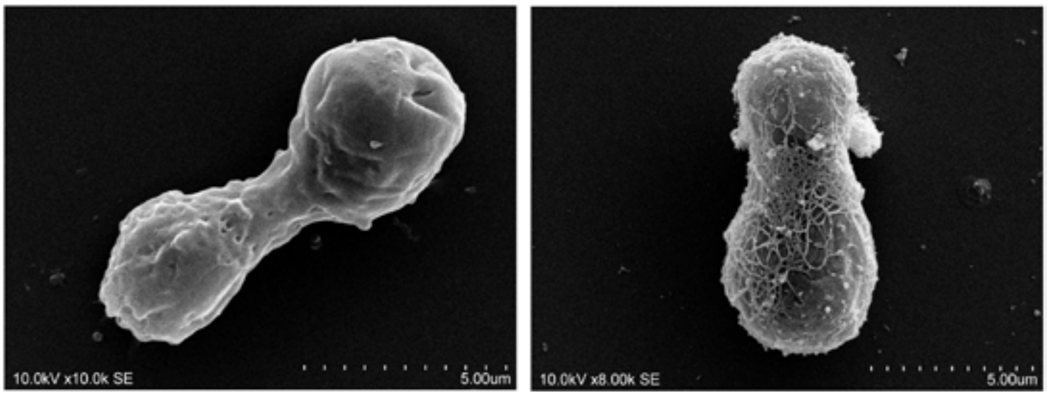

**Supplementary figure 3**

(A) *cps1*-191 *vps20Δ* GFP-*psy1 rlc1*-tdTomato

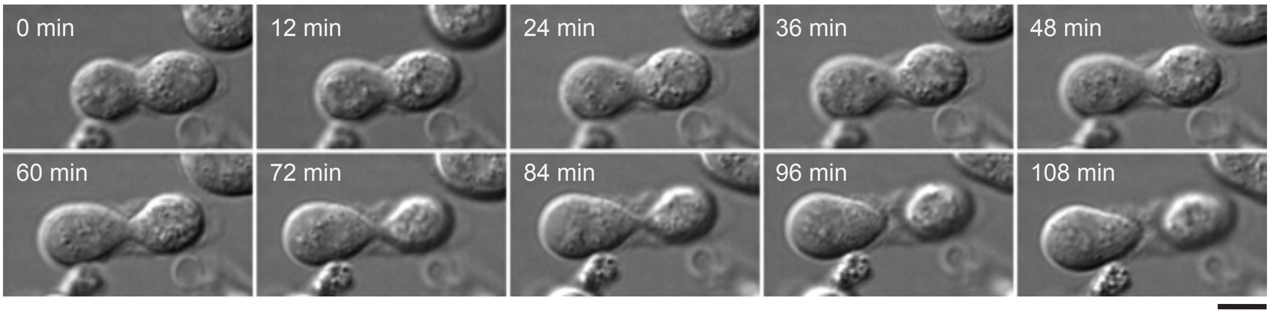


(B) *cps1*-191 *vps4Δ* GFP-*psy1 rlc1*-tdTomato

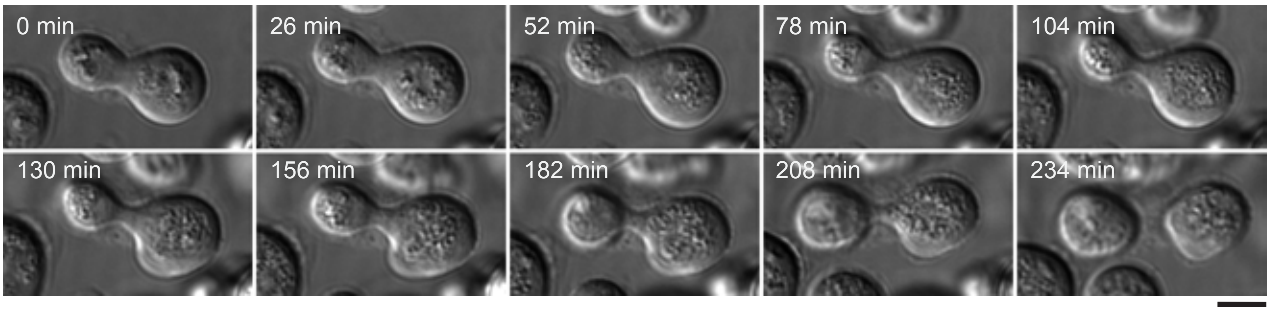

**Supplementary figure 4**

(A) *rlc1Δ cyk3*-GFP cells

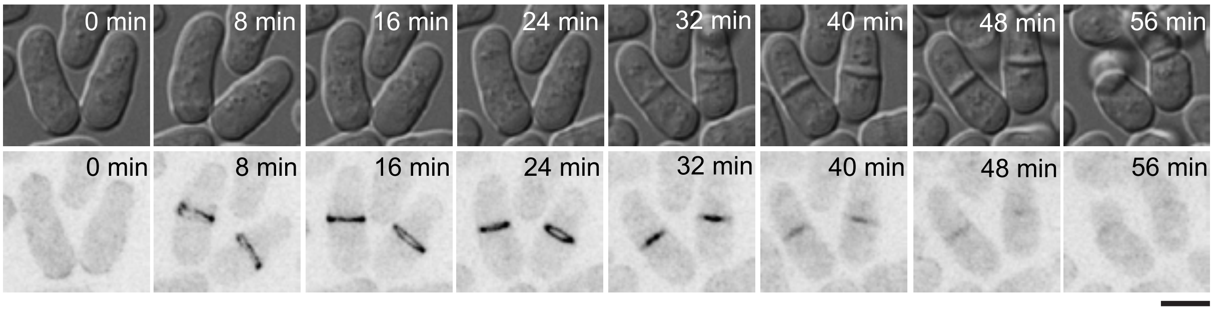
